# High-resolution cryo-EM using beam-image shift at 200 keV

**DOI:** 10.1101/2020.01.21.914507

**Authors:** Jennifer N. Cash, Sarah Kearns, Yilai Li, Michael A. Cianfrocco

## Abstract

Recent advances in single-particle cryo-electron microscopy (cryo-EM) data collection utilizes beam-image shift to improve throughput. Despite implementation on 300 keV cryo-EM instruments, it remains unknown how well beam-image shift data collection affects data quality on 200 keV instruments and how much aberrations can be computationally corrected. To test this, we collected and analyzed a cryo-EM dataset of aldolase at 200 keV using beam-image shift. This analysis shows that beam tilt on the instrument initially limited the resolution of aldolase to 4.9Å. After iterative rounds of aberration correction and particle polishing in RELION, we were able to obtain a 2.8Å structure. This analysis demonstrates that software correction of microscope aberrations can provide a significant improvement in resolution at 200 keV.

## INTRODUCTION

In order to increase the throughput from cryo-EM instruments, many laboratories and facilities have begun using beam-image shift for data collection (Cheng *et al*., 2018). Using this approach, instead of moving the stage to each position on the cryo-EM grid, a process that requires precise movement, the beam is moved in conjunction with image adjustments. Without long waiting times of moving the stage, tilting the beam leads to a dramatic increase in the number of exposures per hour. As such, it is now routine to use beam-tilt to collect 100-300 exposures whereas previously it was only possible to collect 40-50 per hour. This throughput will continue to increase with the advent of direct detectors with faster frame rates, leading to hundreds of exposures per hour.

Even though users can collect 2-3X the amount of data using beam-image shift, they must overcome an additional aberration induced by the beam-image shift: beam tilt (Glaeser *et al*., 2011). When using beam-image shift for collecting exposures, the resulting image will have both axial and off-axis beam tilt (or coma), aberrations that will dampen high-resolution (<3Å) information in the micrographs (Glaeser *et al*., 2011). Due to this, it is a common practice to minimize beam tilt in the cryo-EM instrument through microscope alignments ahead of data collection.

Axial beam tilt aberrations can be corrected computationally for high-resolution structures. For example, this was implemented by Henderson and coworkers for the atomic-resolution structure of bacteriorhodopsin from 2D crystals (Henderson *et al*., 1986). Since its use 40 years ago, recent advances in single-particle cryo-EM have led to the incorporation of axial beam tilt correction into software packages such as RELION (Herzik *et al*., 2017; Wu *et al*.; Zivanov *et al*., 2020). The availability of axial beam tilt correction has led to its widespread adoption for cryo-EM structure determination. Typically, users are finding a 0.2-0.8 mrad beam tilt on previously aligned 300 keV Titan Krios instruments, and correction for this has led to modest improvements in resolution (typically 0.1 - 0.3Å) (Zivanov *et al*., 2018).

Even though beam-image shift data collection in combination with aberration correction has been implemented for datasets at 300 keV, there is limited information on how much beam tilt is induced by beam-image shift at 200 keV and if it can be overcome computationally. Given that the phase error caused by either axial or off-axis beam tilt scales with the wavelength (λ) squared (Glaeser *et al*., 2011), changing from 300 keV (λ = 1.96 pm) to 200 keV (λ = 2.51 pm) will result in worse phase error from both axial and off-axis beam tilt. While previous work indicated that short-range beam-image shift could achieve a 3.3Å for the T20S proteasome at 200 keV (Herzik *et al*., 2017), this same work required using stage position to obtain a resolution better than 3Å. Recently, using these original datasets of aldolase and T20S datasets, RELION-3.1 now allow higher-order aberrations to be corrected computationally (Zivanov *et al*., 2020; Wu *et al*.). This allowed the resolution of aldolase to improve from 2.5Å to 2.1Å and the T20S proteasome improved from 3.1Å to 2.3Å.

In order to test the limits of computational correction of microscope aberrations at 200 keV, we collected and analyzed a dataset of aldolase using beam-image shift on a Talos-Arctica at 200 keV. Using this dataset, we were able to determine a 4.9Å structure of aldolase without aberration corrections. Following iterative rounds of axial beam tilt correction and particle polishing, we were able to determine a 2.8Å structure of aldolase. This indicates that beam-image shift can be an effective data collection strategy to increase the throughput on 200 keV cryo-EM instruments, where microscope aberrations can be corrected computationally.

## RESULTS

### Beam-image shift data collection & analysis

In order to test the impact of beam-image shift on data quality, we set up the automated data collection system to target 5×5 areas with beam-image shift (**Figure 1A**). At medium magnification (**Figure 1A**), we typically focused on the middle hole which was followed by beam-image shift with distances up to 5 μm away from the beam center. After collecting 2,111 micrographs, we obtained a large range of beam-image shift micrographs that provided a near-continuous distribution across the 10 x 10 μm area (**Figure 1B**). Interestingly, while many micrographs showed minimal objective astigmatism (**Figure 2A, left**), a large percentage of the dataset showed exaggerated objective astigmatism (**Figure 2A, right**) which can be induced by a large amount of axial beam tilt (Glaeser *et al*., 2011).

**Figure 1.**
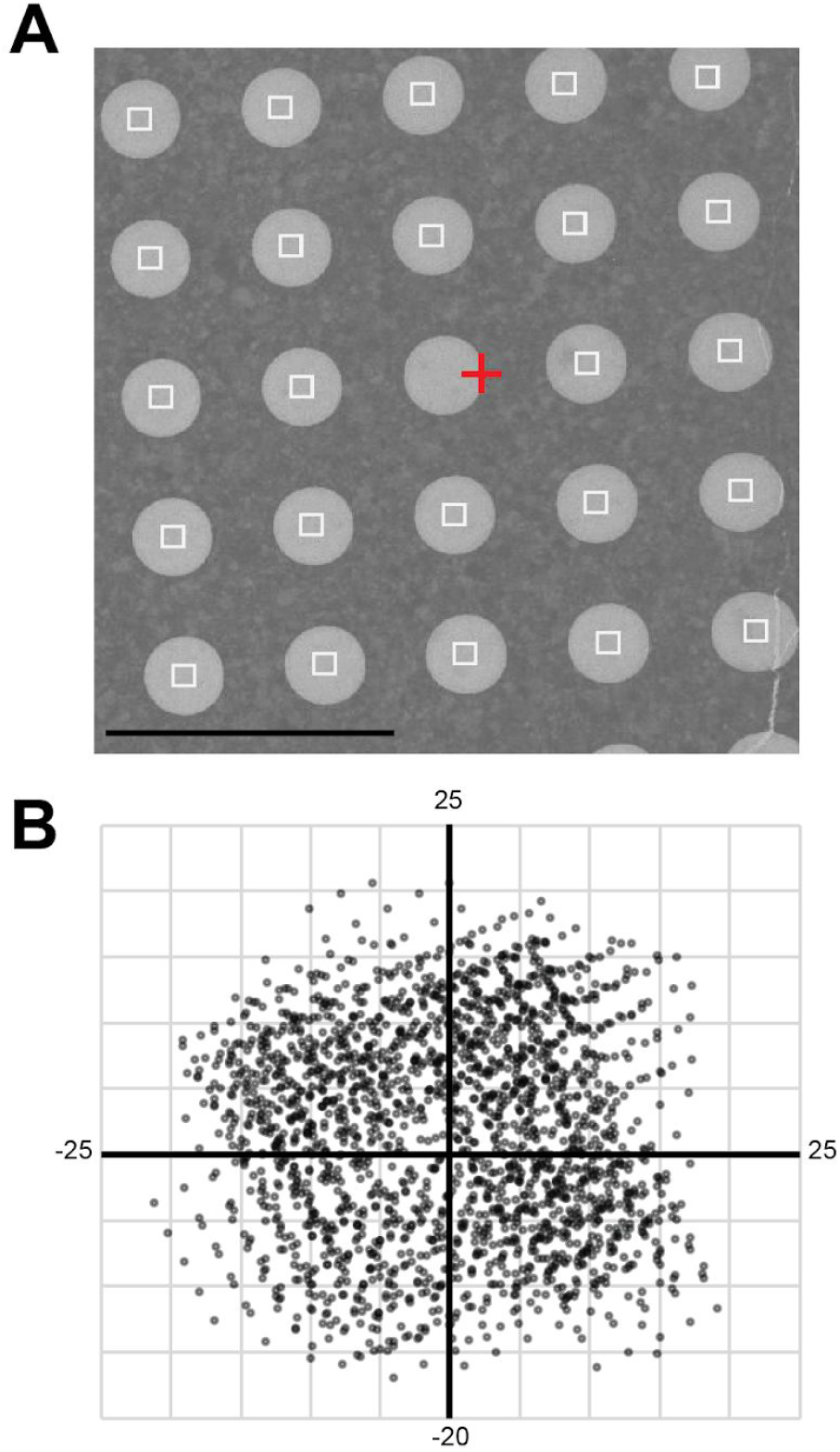
Data collection strategy tor micrographs collected with beam-image shift. (A) Representative image at intermediate magnification. Red cross: focus area; White squares: exposures; Scale bar is 5 μm. Each exposure was collected with image-shift beam tilt. (B) Overview of image shift values from Leginon for beam tilt dataset. Units shown are μm.

**Figure 2.**
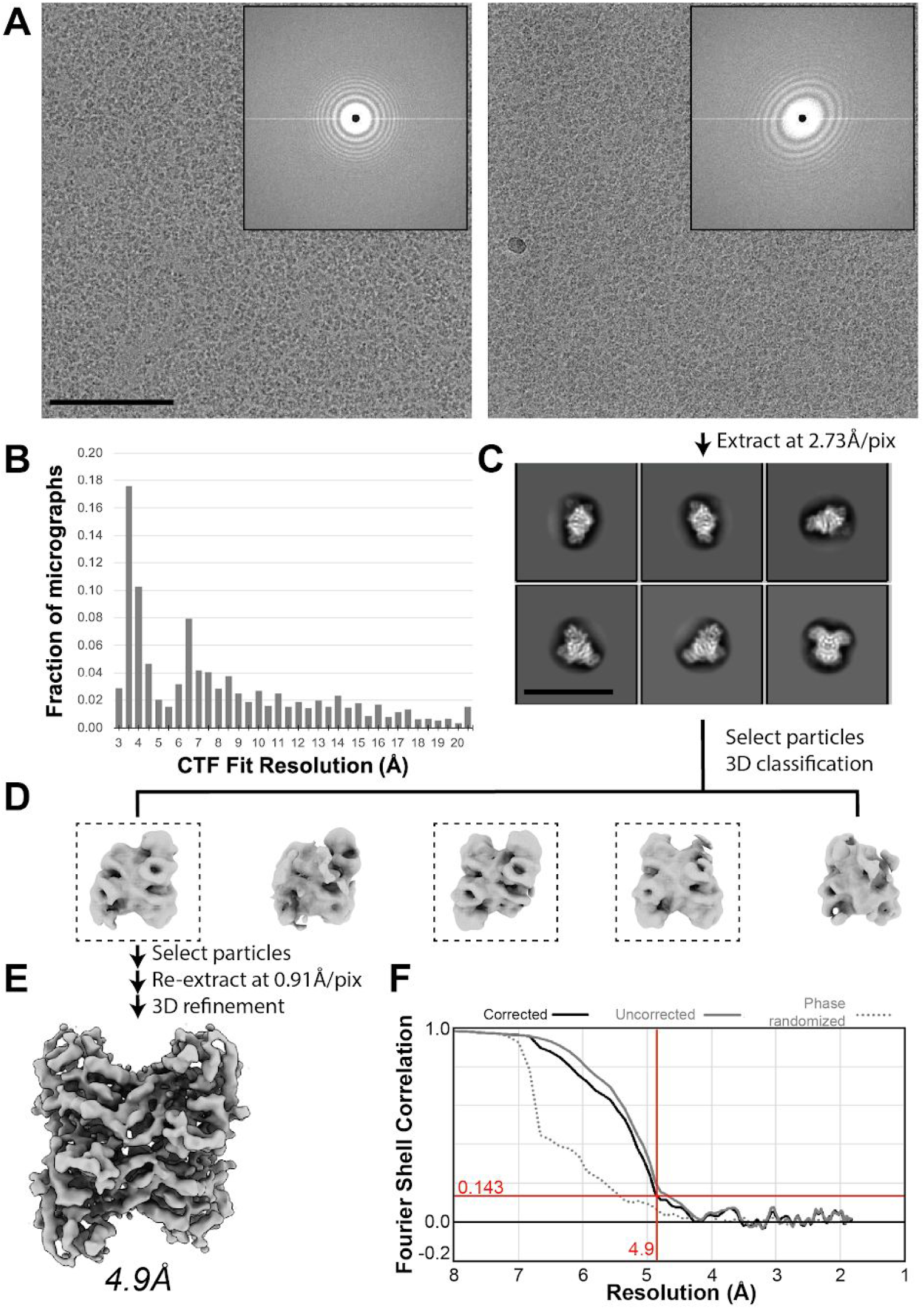
Single particie analysis of aldolase without beam-tilt correction. **(A)** Representative micrographs with minimal (left) and obvious (left) beam tilt-induced objective astigmatism inset: Cropped power spectrum. Scale bar is 100 nm. (B) Histogram of CTF resolution limits across dataset using CTFFIND4. (C) Representative 2D class averages calculated using RELION. Scale bar is 200Å, (D) 3D classification resuits for selected particles after 2D classification. Dashed boxes indicate classes with particles used for subsequent 3D refinement. (E) Sharpened reconstruction after 3D refinement using RELION filtered to 4.9Å. (F) FSC curves for final reconstruction.

Following data collection, the aldolase beam-image shift data were analyzed using standard single-particle processing (**Figure 2**). This involved estimating the contrast transfer function (CTF) using CTFFIND4 (Rohou & Grigorieff, 2015), which yielded CTF fits to higher than 4Å resolution for the majority of the micrographs (**Figure 2B**). After picking and extracting particles, 2D classification showed clear secondary structure features (**Figure 2C**), consistent with previous work on aldolase (Herzik *et al*., 2017; Kim *et al*., 2018). After selecting particles from class averages exhibiting high-resolution features, we performed 3D classification in order to obtain a homogenous population of aldolase particles with all four subunits intact (**Figure 2D**). Using these selected particle coordinates, particles were re-extracted at the full pixel size (0.91 Å/pixel) and subjected to 3D refinement in RELION. The refined structure reached a resolution of only 4.9Å (**Figure 2E & 2F**), which is significantly less than published work of ~3Å (Kim *et al*., 2018; Herzik *et al*., 2017). This suggested that the aberrations from beam tilt induced by beam-image shift data collection are likely limiting the resolution of the final structure.

### Beam tilt correction of aldolase cryo-EM micrographs

After determining a refined 3D structure of aldolase, we wanted to test whether the beam tilt refinement option in RELION 3.0 is capable of overcoming such a large degree of axial beam tilt. To use this feature of RELION, the micrographs must be grouped into beam tilt groups. Considering the near-continuously changing beam-image shift data collection for the entire dataset (**Figure 1B**), beam-image shift values from Leginon were used in order to divide the micrographs into groups (**Figure 3A**). This involved dividing data into groups of 25 (5×5), 100 (10×10), and 400 (20×20) based on the amount of Resolution (A) beam-image shift in Leginon (**Supplemental Figure 1**). For each grouping, the particles underwent beam tilt refinement, 3D refinement, and sharpening in RELION in order to determine the change in the final resolution of the structure. We saw that grouping into 5×5, 10×10, and 20×20 groups had a significant increase in the final resolution of 4.1Å, 4.0Å, and 3.8Å, respectively (**Figure 3B**). This result indicates that the previously determined structure at 4.9Å was limited in resolution due to beam tilt aberrations that could be partially overcome by grouping the data into beam tilt groups in RELION.

**Figure 3.**
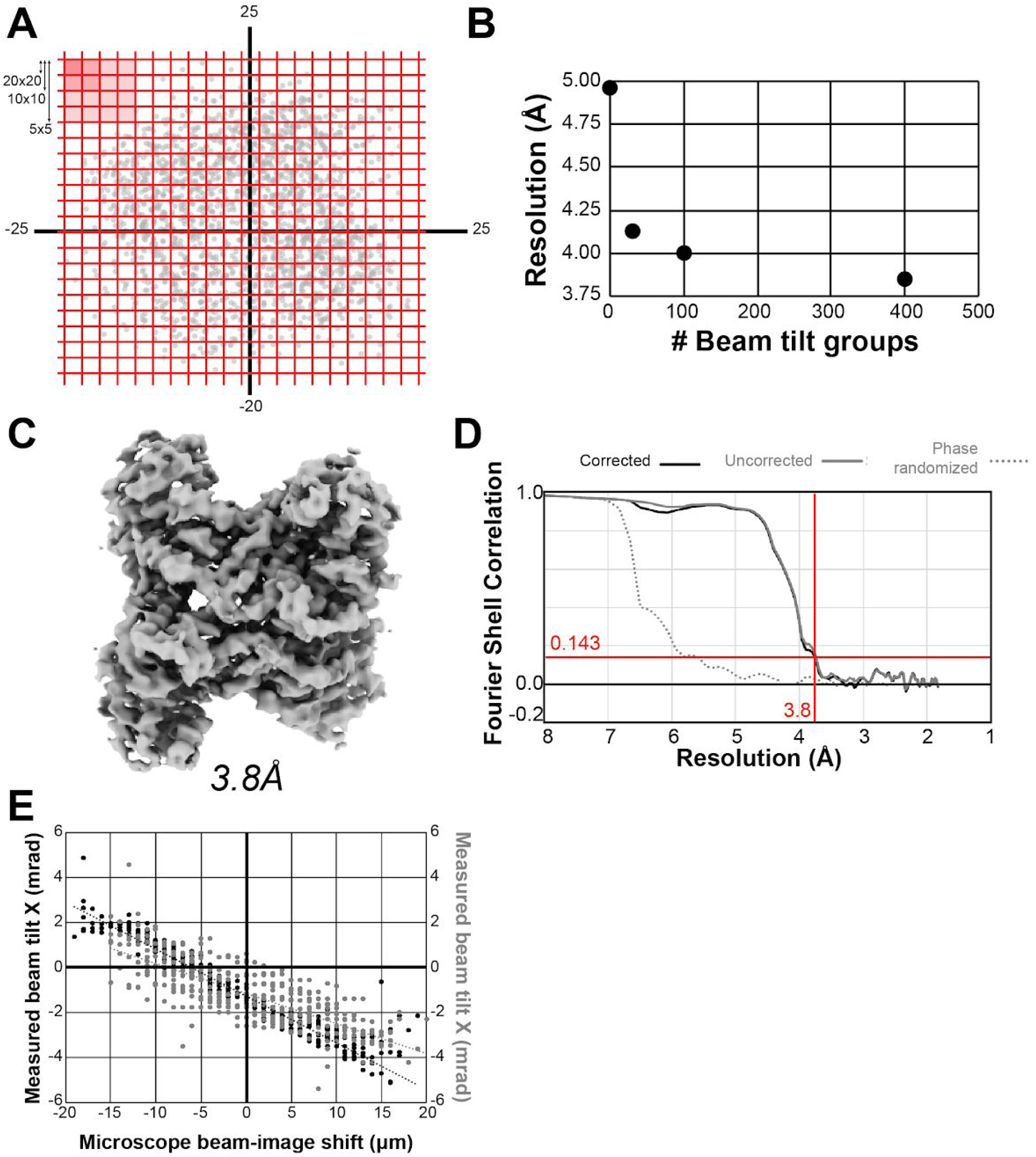
Improved resolution and map quality using beam tilt refinement. **(A)** Strategy for grouping micrographs. Micrographs were grouped into 25 groups (5×5), 100 groups (10×10), and 400 groups (20×20). (B) Effect of group size on beam tilt refinement and subsequent resolution estimation for refined 3D structures. (C) Sharpened 3D reconstruction for particles places into 400 micrograph groups filtered to 3.8Å. (D) FSC curves for 3D reconstruction in (C). (E) Beam tilt measurements for each group displayed with respect to microscope beam-image shift for X (black) and Y coordinates (gray). Dashed lines show least squares fit where R^2^=0.96 (beam tilt X) and R^2^=0.64 (beam tilt Y).

For the micrographs divided into 400 groups, the subsequently refined map showed improved density features and had a gold standard FSC value of 3.8Å (**Figure 3C & 3D**). This indicates that beam tilt refinement improved the resolution of aldolase significantly from 4.9Å to 3.8Å in a single step.

Using the calculated beam tilt values from RELION, we then compared how beam tilt changed as a function of microscope beam-image shift (**Figure 3E**). This comparison reveals a few key features of this dataset. First, without any applied beam-image shift at [0,0], there was a significant amount of beam tilt present: −1.24 mrad (X) and −1.14 mrad (Y). Second, the change in beam tilt based on change in beam-image shift (the slope in **Figure 3E**) was different for the X versus Y direction: −2.1e05 μm/mrad vs. −1.35e5 μm/mrad, respectively. Finally, this result also shows that a subset of micrographs have a much larger beam tilt than the majority of micrographs, explaining why some micrographs displayed objective astigmatism due to high beam tilt (**Figure 2C**).

Given that the RELION beam tilt estimation step is dependent on the resolution of the 3D reconstruction, we performed iterative beam tilt refinements and Bayesian particle polishing in order to test whether refinement of beam tilt and particles can further increase the dataset resolution (**Figure 4**). Starting with the 20×20 grouped dataset at 3.8Å reconstruction (**Figure 4B**), we used this map to re-calculate beam tilt for micrographs across the dataset. Then, using these new beam tilt values, we performed another round of 3D refinement. This new structure refined to higher resolution at 3.6Å and had a lower B-Factor (−105Å^2^) (**Figure 4C**), indicating per-particle quality has increased. After these two rounds of beam-tilt refinement, we then utilized Bayesian particle polishing in RELION (Zivanov *et al*., 2019) to further improve the resolution to 3.3Å (B-Factor −91Å^2^) (**Figure 4D**). Then, with these particles, we performed a final beam tilt calculation, allowing us to determine a 2.8Å reconstruction (B-Factor −52Å^2^) (**Figure 4E**). This reconstruction could not be improved with further aberration refinements or defocus refinements (data not shown). The increase in map quality and model statistics from 4.9Å to 2.8Å (**Supplemental Figure 2, Supplemental Table 2**) demonstrates that the aberration correction improved the interpretability of the reconstructions.

**Figure 4.**
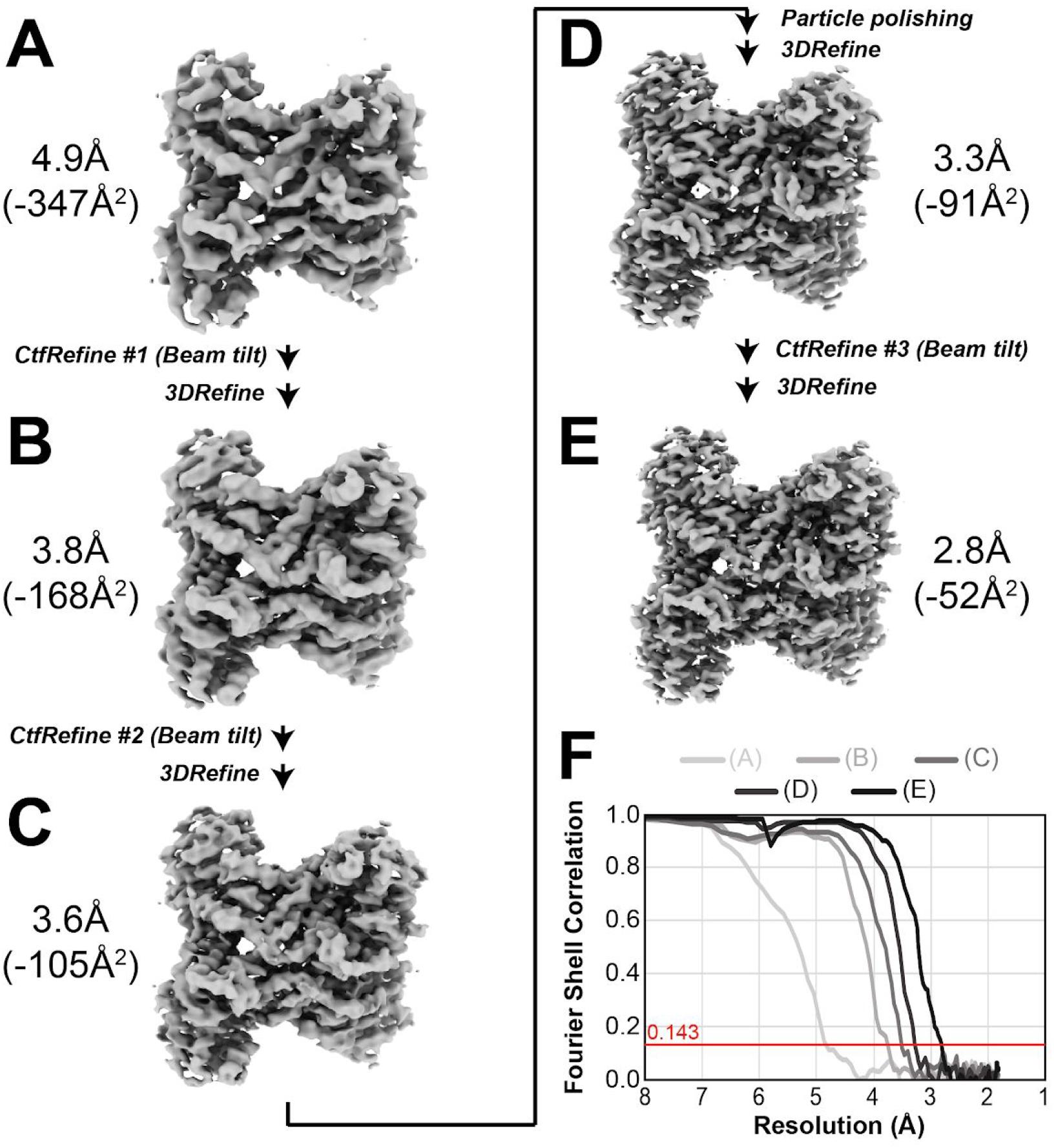
Iterative CTF-refinement with particle polishing improves overall resolution to 2.8Å. (A) Initial 3D structure at 4.9Å. Following the first CTF refinement and 3D refinement to obtain a structure at 3.8Å (B), continued CTF refinements alongside Bayesian particle polishing allowed for resolution and B-factor improvements (C) - (E), ultimately allowing the determination of a 2.8Å structure (E). (F) FSC curves for 3D reconstructions from (A)-(E).

In order to test whether there were remaining beam tilt aberrations, we divided the final reconstruction into two subsets: 1) particles with <0.5 mrad measured beam tilt and 2) particles with >2 mrad measured beam tilt (**Supplemental Figure 3**). After matching the number of particles per group to be the same (group #1 only had 20,231 particles), we refined these two groups using RELION. Group #1 refined to higher resolution and lower B-Factor (3.2Å, −24Å^2^) (**Supplemental Figure 3B**) vs. group #2 (3.5Å, −55Å^2^) **Supplemental Figure 3C**). This indicates that the data quality for the small measured beam tilt group is higher than for particles with larger beam tilt.

The final structure at 2.8Å (**Figure 5**) shows dramatically improved density features compared to the original 4.9Å structure. Specifically, the significantly higher resolution provides unambiguous secondary structure tracing whereas the 4.9Å structure contained many more ambiguities (**Figure 5B**). A comparison of model refinement statistics also highlights the improved map quality for the final 2.8Å reconstruction (**Supplemental Table 2**). This structure demonstrates that computational correction of microscope aberrations and particle motion allows for sub-3Å structure determination.

**Figure 5.**
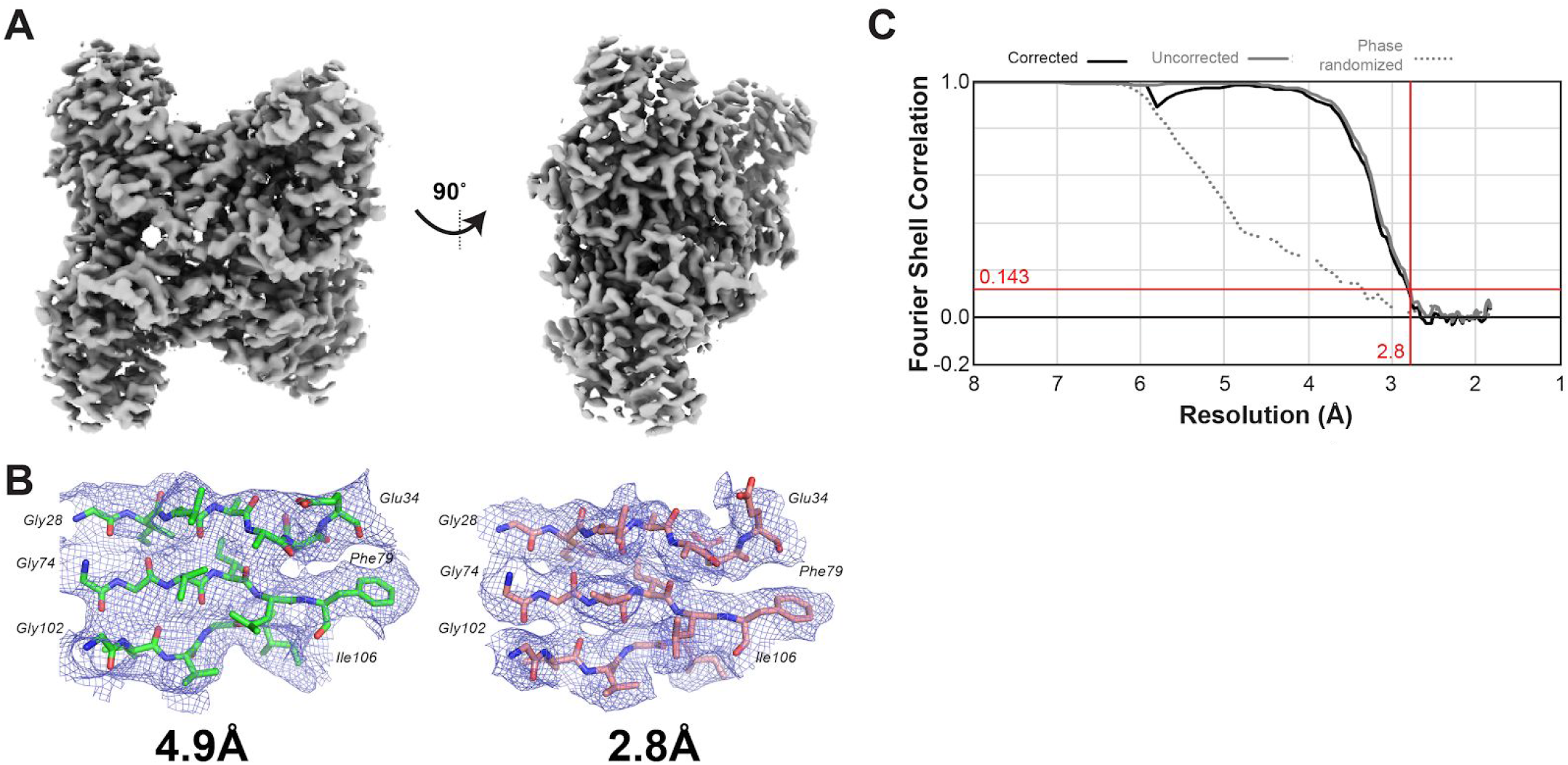
Structure of aldolase at 2,8Å after beam tilt and particle polishing. (A) Sharpened aldolase reconstruction at 2.8Å. (B) Example densities and models for aldolase at 4.9Å and 2.8Å. (C) FSC curves for final reconstruction.

## DISCUSSION

### Single-particle analysis of aldolase with significant microscope aberrations

The dataset analyzed in this work utilized significant beam-image shift data collection at 200 keV on a Talos Arctica. This strategy introduced significant microscope aberrations into the raw data and was significant enough to cause objective astigmatism in micrographs due to a large amount of beam tilt (**Figure 2A, right**).

Despite the presence of significant aberrations, analysis of resulting aldolase particle stacks allowed for 2D and 3D averaging. The 2D class averages obtained from RELION for aldolase (**Figure 2C**) are indistinguishable from previously published aldolase class averages (Herzik *et al*., 2017; Kim *et al*., 2018), indicating that the aberrations do not affect 7-10Å-resolution class averages. Importantly, however, 3D refinement of the original particle stack does not achieve better than 4.9Å resolution (**Figure 2E**), which is much lower than typical aldolase reconstructions that are within the range of 3-4Å for initial 3D refinements (Herzik *et al*., 2017; Kim *et al*., 2018). This analysis indicates that microscope aberrations do not affect sample screening and initial 2D averaging, however, the aberrations prevent structure determination <5Å.

### Significant improvement of resolution through iterative beam tilt correction

By taking advantage of microscope aberration correction in RELION-3.1 (Wu *et al*.; Zivanov et al., 2020) we were able to improve the resolution of aldolase from 4.9Å to 2.8Å. While previous work demonstrated that aberration refinement allows for resolution improvements for data at both 300 keV (Zivanov *et al*., 2018) and 200 keV (Zivanov *et al*., 2020; Wu *et al*) all datasets analyzed were collected on relatively well-aligned instruments. With high-quality starting data, the initial reconstructions prior to aberration correction achieved ~3Å (unlike this work which was 4.9Å). Moreover, the data collected at 200 keV (Wu et al.; Herzik *et al*., 2017) used stage position instead of beam-image shift, further minimizing microscope aberrations in the dataset.

Using these algorithmic improvements (Zivanov *et al*., 2018) in combination with Bayesian particle polishing (Zivanov *et al*., 2019), we were able to improve the resolution of aldolase to 2.8Å (**Figure 4 & 5**). Analysis of the measured beam tilts indicates that there was axial beam tilt present on the instrument prior to using beam-image shift (**Figure 3E**). This confirms that the microscope had axial beam tilt prior to data collection, where better microscope alignments could have minimized this issue.

Despite utilizing microscope aberration correction and particle polishing, the overall per-particle data quality remains worse than stage position-collected aldolase data. By comparing the final B-Factor from our data collected using beam-image shift (−52Å^2^) with aldolase determined from stage position (−35Å^2^) (Herzik *et al*., 2017), the higher B-Factor for our data indicates that per-particle signal is lower for our dataset. Importantly, for particles with <0.5 mrad beam tilt, we obtained a B-factor of −24Å^2^, indicating that a subset of particles was of comparable or higher quality than published work. We do not know if alternative data processing strategies are needed for beam-image shift data collection or whether our sample preparation of aldolase is of poorer quality, but further work is needed to verify if beam-image shift B-Factors are consistently higher than stage position collected data at 200 keV.

### Data throughput vs. data quality

The main motivation to utilize beam-image shift for data collection instead of stage position is the increased data collection throughput. For the dataset collected here, we were able to obtain a 2.4X increase in throughput for beam-image shift when compared with stage position: 73 movies per hour (beam-image shift) vs. 30 movies per hour (stage position). Considering the cost of instrument time, beam-image shift provides 1,752 movies per 24 hour period vs. 720 movies per 24 hour period for stage position. Indeed, the latest generation of detectors that have faster readout stands to triple this throughput for beam-image shift.

Based on our analysis of aldolase, we believe that there is a significant difference between 200 keV vs. 300 keV beam-image shift data collection (for instances where there is not an optical correction on the microscope). At 300 keV, it is possible to use a comparable beam-image shift as that used in this study but instead obtain a structure ~3Å (Zivanov *et al*., 2018). For this dataset at 300 keV, beam-image shift provides high-resolution structures prior to aberration correction. Unlike this previous study, the aldolase structure collected using beam-image shift at 200 keV was limited in resolution due to aberrations to 4.9Å. In order to correct for the aberrations, significant effort was required in order to perform optical grouping and analysis, steps that may be beyond beginning to intermediate RELION users.

With these considerations, we advocate beam-image shift at 200 keV for sample screening. This is because we observed high-quality 2D class averages for aldolase despite significant beam tilt, information well-suited for sample screening (i.e. changing buffers, sample concentrations, etc.). However, this study does indicate that even if a user collected data with significant beam tilt from beam-image shift data, software-based aberration correction is possible to <3Å for well-behaved samples like aldolase.

## DATA ACCESSIBILITY

Cryo-EM structures have been deposited to the EMDB under accession codes EMDXXXX, .… All movies, micrographs, particle stacks, and metadata files are deposited to EMPIAR under XXXX.

## ACKNOWLEDGMENTS

We would like to thank all members of the cryo-EM community at the University of Michigan. We would like to particularly thank Dr. Takanori Nakane, Wim Hagen, & Dr. Min Su for feedback on this work. This work was supported by NSF-DBI-ABI 1759826 (Y.L. & M.A.C.). The research reported in this publication was supported by the NIH under award number S10OD020011.

## METHODS

### Sample preparation

Pure aldolase isolated from rabbit muscle was purchased as a lyophilized powder (Sigma Aldrich) and solubilized in 20 mM HEPES (pH 7.5), 50 mM NaCl at 1.6 mg/ml. Sample as dispensed on freshly plasma cleaned UltrAuFoil R1.2/1.3 300-mesh grids (Electron Microscopy Services) and applied to grid in the chamber of a Vitrobot (Thermo Fisher) at ~95% relative humidity, 4°C. Sample was blotted for 4 seconds with Whatman No. #1 filter paper immediately prior to plunge freezing in liquid ethane cooled by liquid nitrogen.

### Cryo-EM data acquisition and image processing

Data were acquired using the Leginon automated data-acquisition program (Suloway *et al*., 2005). Image pre-processing (frame alignment with MotionCor2 (Zheng *et al*., 2017) and CTF estimation using CTFFIND4 (Rohou & Grigorieff, 2015)) were done using the Appion processing environment (Lander *et al*., 2009) for real-time feedback during data collection. Images were collected on a Talos Arctica transmission electron microscope (Thermo Fisher) operating at 200 keV with a gun lens of 6, a spot size of 6, 70 μm C2 aperture and 100 μm objective aperture using beam-image shift. Movies were collected using a K2 direct electron detector (Gatan Inc.) operating in counting mode at 45,000x corresponding to a physical pixel size of 0.91 Å/pixel with a 10 sec exposure using 200 ms per frame. Using an exposure rate of 4.204 e/pix/sec, each movie had a total dose of approximately 42 e/Å^2^ for the 2,111 movies over a defocus 0.8-2 μm.

### Pre-processing

Movies were aligned using RELION-3.0 (Zivanov *et al*., 2018) (3.0-beta-2) motion correction with 5 patches in both X & Y directions, a B-Factor of 150Å^2^ without binning. Following motion correction, CTF estimation was performed with CTFFIND4 (Rohou & Grigorieff, 2015) using exhaustive search for a defocus range of 0.5 to 5.0 μm (0.05 μm step size) and an astigmatism search range of 0.5 μm within a resolution range of 6 and 30Å. The combination of a large astigmatism search with exhaustive searches led to many over-estimates of CTF resolution fits for this dataset. Therefore, in order to remove micrographs automatically, we utilized our recently developed *MicAssess (Li et al.)* program to remove all empty and bad micrographs. This removed 685 micrographs, leaving 1,426 micrographs for particle picking. Particles were picked from aligned micrographs using crYOLO (Wagner *et al*., 2019) general model PhosaurusNet with an anchor size of 98 x 98 pixels.

### Single-particle analysis without aberration correction

For 2D classification, 718,578 particles were extracted with an unbinned box size of 300 pixels and subsequently binned to 2.73Å (box size 100 pixels). Particles were then subjected to 2D classification into 100 classes using RELION-3.0.2 (T=2; Iter=25). After selecting particles from the best classes, 275,487 particles underwent 3D classification into 5 classes using RELION-3.0.2 (T=4; Iter=25) and EMD-8743 (Herzik *et al*., 2017) as a reference model. Following the selection of the best classes, 186,841 particles were centered and re-extracted at 0.91Å/pixel. This stack was used for 3D refinement to obtain a post-processed structure with a resolution of 4.9Å and a B-Factor of −347Å^2^.

### Aberration correction and particle polishing

Particles were grouped into optics groups based on beam-image shift values obtained from the Leginon database. In order to group particles into discrete optics groups, the entire file of beam-image shift values were divided into 5×5, 10×10, or 20×20 groups. The first two beam tilt estimation steps (CtfRefine #1 & #2, Figure 4) used RELION-3.0 (3.0-beta-2). Subsequent steps (Bayesian polishing and CtfRefine #3) used RELION-3.1 (version 30001). All steps for aberration correction and polishing are described in Figure 4. Aberration correction and polishing did not improve resolution more than the final 2.8Å aldolase structure. We also tested whether using predicted beam tilts from CtfRefine #1 could improve the resolution of a final reconstruction, however, this did not improve dataset resolution (data not shown).

### Model building and refinement

The coordinates for rabbit aldolase (PDB: 5vy5) were docked into each map in PHENIX using phenix.dock_in_map (Adams *et al*., 2012). Structure refinement and model validation were performed using phenix.real_space_refine (Afonine *et al*., 2018). The same docking and refinement parameters were used for each map. To make figures showing map density, phenix.map_box was used to restrict the map shown to specific stretches of residues. Root mean square deviation (rmsd) values comparing all atoms between structures were calculated using a Least Squares Fit in Coot (Emsley *et al*., 2010). The PyMOL Molecular Graphics System (Version 2.1, Schrödinger, LLC) was used to render images showing these structures.

## Supplemental materials

**Supplemental Figure 1.**
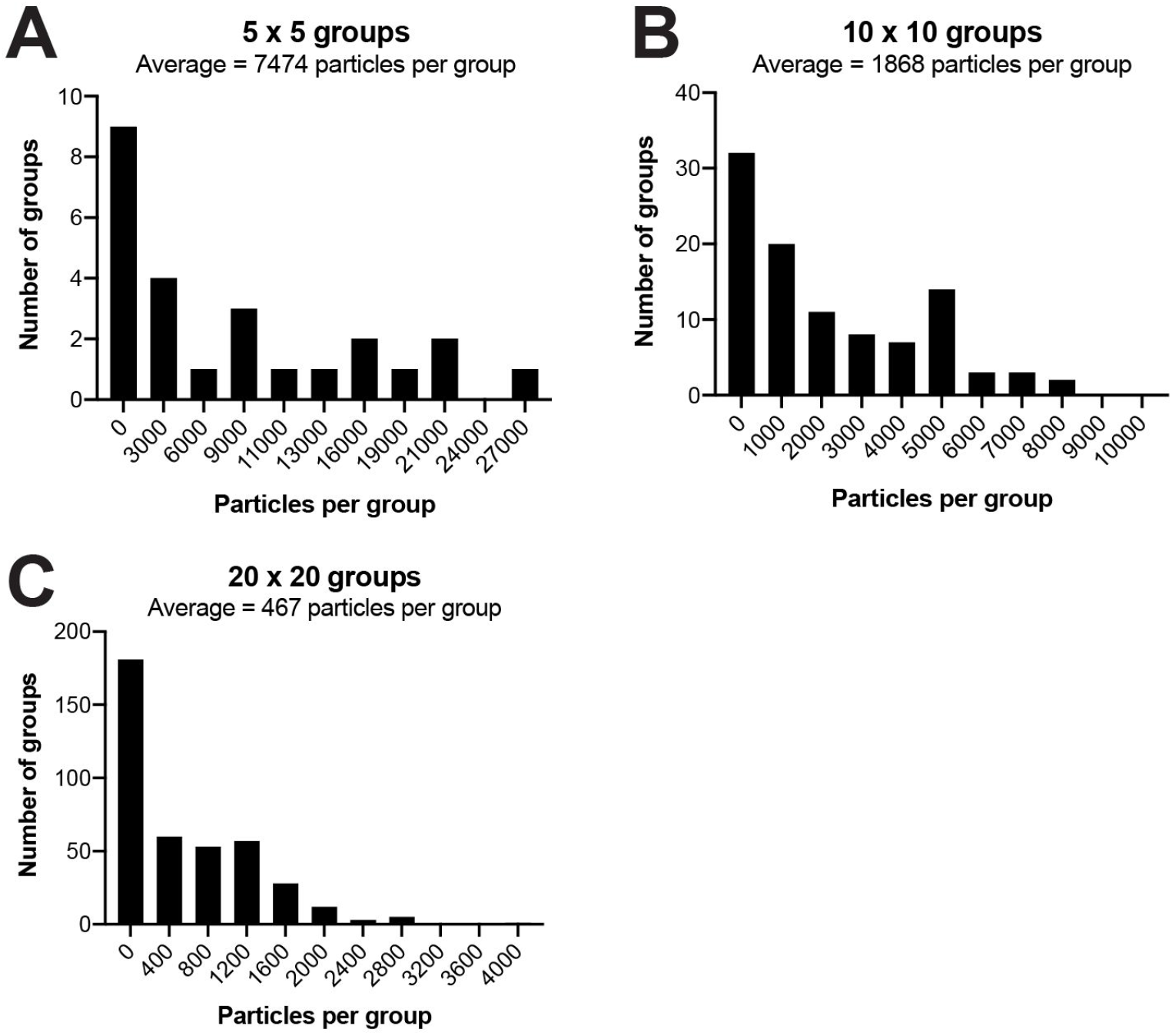
Particle numbers per group. Histograms of particle distribution per beam tilt grouping. (A) 5×5, (B) 10×10, (C) 20×20.

**Supplemental Figure 2.**
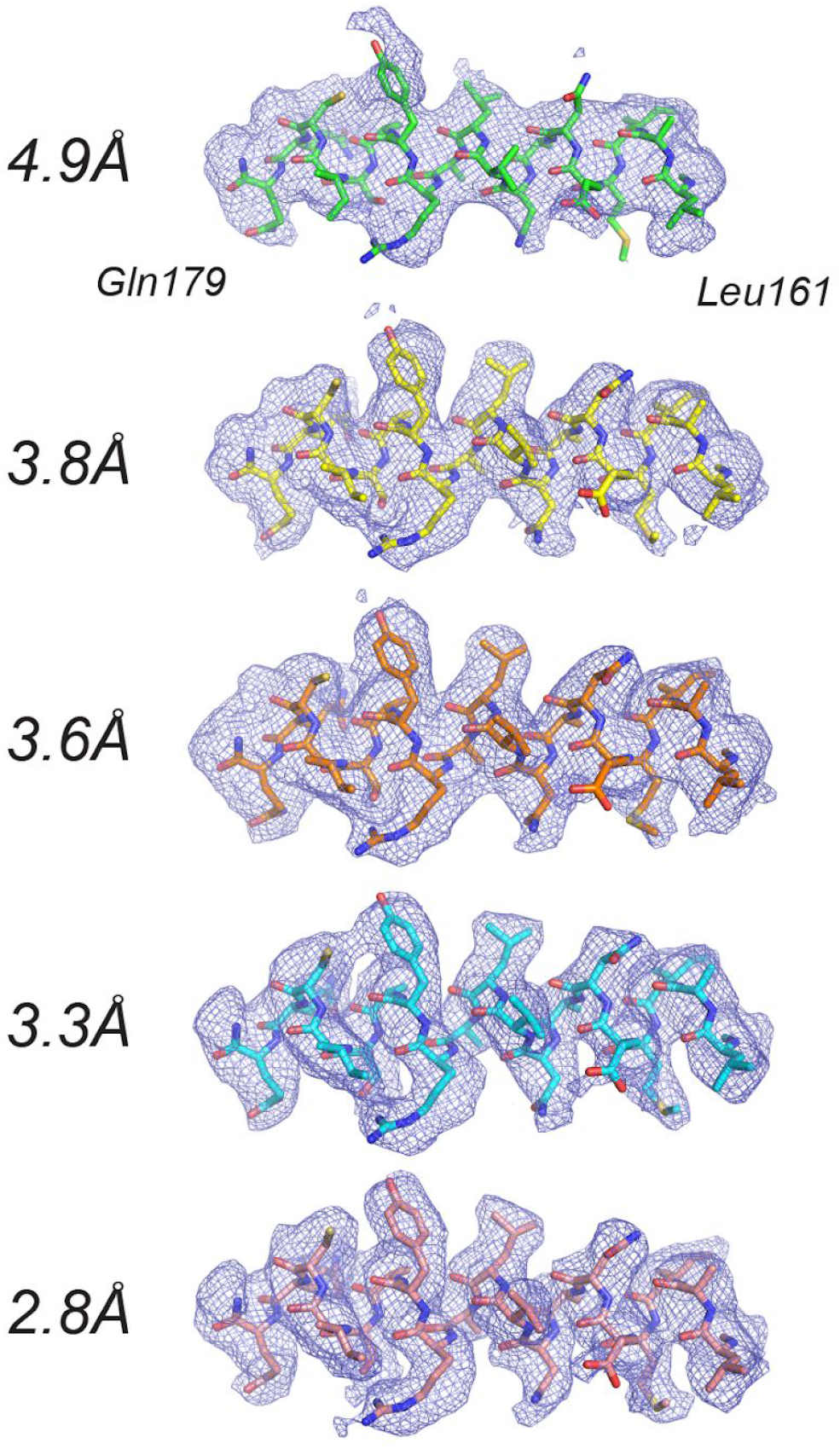
Representative densities from iterative beam-tilt refinements. Sharpened densities with associated models highlight changes in density quality through iterative rounds of beam-tilt refinement corresponding to structures from Figure 4A-4E.

**Supplemental Figure 3.**
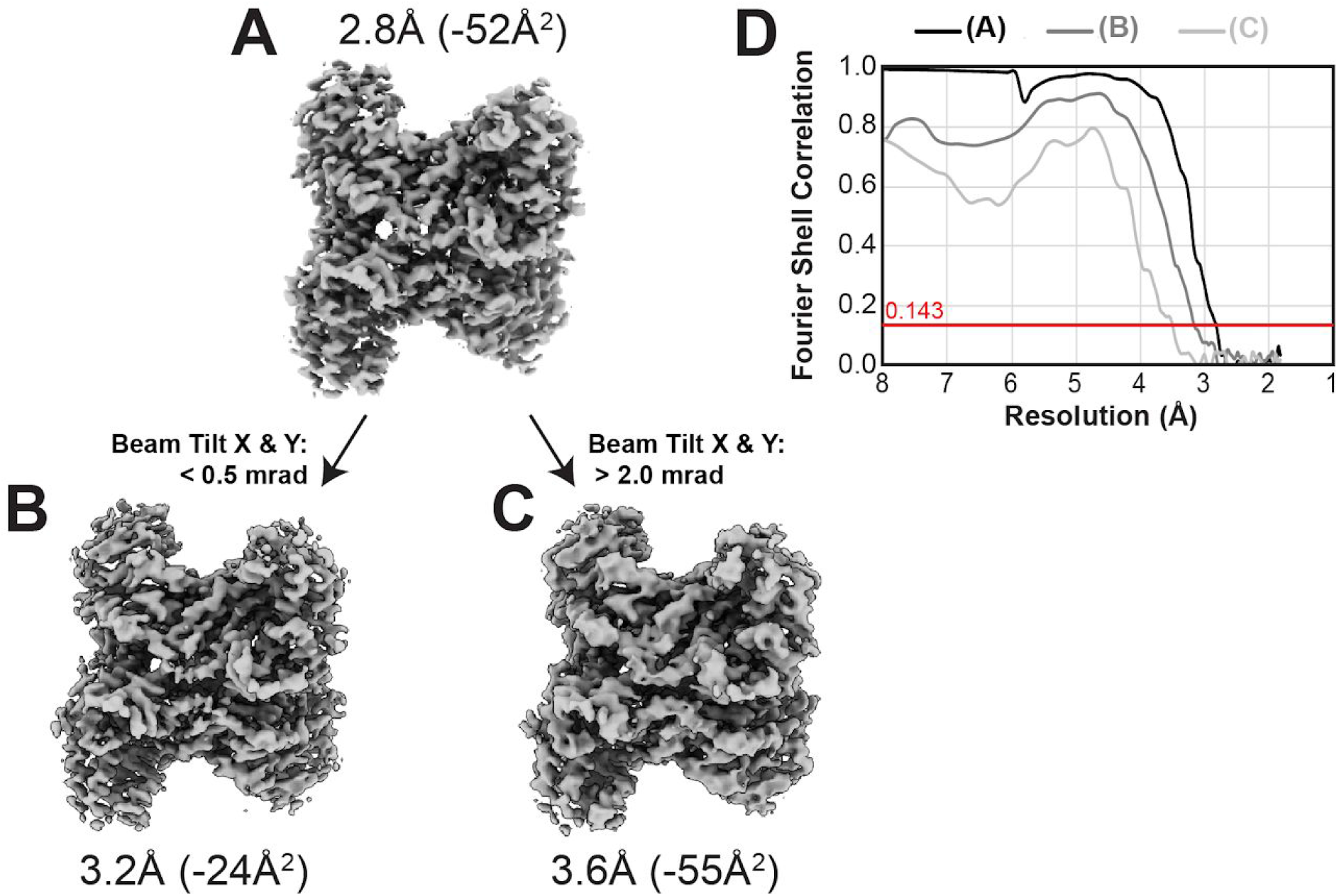
Re-analysis of beam tilt subgroups from final reconstruction. (A) Final reconstruction of aldolase at 2.8Å. (B) Reconstruction of aldolase from micrographs with < 0.5 mrad beam tilt at 3.2Å. (C) Reconstruction of aldolase from micrographs with > 2.0 mrad beam tilt at 3.5Å. (D) FSC curves.

**Supplemental Table 1.** Cryo-EM data collection, refinement and validation statistics.

|  | **Aldolase (19apr12a)** |
| --- | --- |
| **Microscope** | Talos Arctica |
| **Detector** | Gatan K2 |
| **Voltage (kV)** | 200 |
| **Electron exposure (e<sup>-</sup>/Å<sup>2</sup>)** | 43 |
| **Defocus range (μm)** | 0.8 - 2 |
| **Data collection mode** | Beam-image shift |
| **Micrographs collected (per hour)** | 73 |
| **Original pixel size (Å)** | 0.91 |
| **Symmetry imposed** | D2 |
| **Initial number of micrographs** | 2,111 |
| **Final number of micrographs** | 569 |
| **Initial particle images (no.)** | 718,578 |
| **Final pixel size** | 0.91 |
| **Final particle images (no.)** | 186,841 |
| **Number of optics groups** | 400 |
| **FSC threshold** | 0.143 |
| **Final map resolution (Å)** | 2.8 |
| **Final B-Factor (Å<sup>2</sup>)** | -52 |

**Supplemental Table 2.** Model building statistics.

| **Structure from Figure 4** | **4A** | **4B** | **4C** | **4D** | **4E** |
| --- | --- | --- | --- | --- | --- |
| **Resolution (Å)** | 4.9 | 3.8 | 3.6 | 3.3 | 2.8 |
| **B-Factor (Å<sup>2</sup>)** | -347 | -168 | -105 | -91 | -52 |
| **Bonds (RMSD)** |  |  |  |  |  |
| Length (Å) | 0.007 | 0.008 | 0.008 | 0.004 | 0.005 |
| Angles (°) | 1.197 | 0.970 | 0.973 | 0.767 | 0.894 |
| **Molprobity score** | 2.18 | 2.21 | 1.94 | 1.35 | 1.86 |
| **Clash score** | 9.26 | 6.36 | 6.08 | 3.99 | 4.23 |
| **Ramachandran plot (%)** |  |  |  |  |  |
| Outliers | 0 | 0 | 0 | 0 | 0 |
| Allowed | 3.52 | 3.81 | 4.40 | 2.93 | 3.23 |
| Favored | 96.48 | 96.19 | 95.60 | 97.07 | 96.77 |
| **Rotamer outliers (%)** | 3.96 | 6.12 | 2.52 | 0.36 | 3.96 |
| **CaBLAM outliers (%)** | 3.54 | 2.95 | 2.95 | 2.95 | 2.65 |
| **CC (mask)** | 0.78 | 0.81 | 0.83 | 0.82 | 0.82 |
| **CC (box)** | 0.67 | 0.79 | 0.87 | 0.76 | 0.77 |
| **CC (peaks)** | 0.58 | 0.71 | 0.80 | 0.72 | 0.73 |
| **CC (volume)** | 0.79 | 0.81 | 0.83 | 0.81 | 0.81 |
| **RMSD (Å) all atoms compared to Figure 4E** | 1.01 | 0.80 | 0.84 | 0.62 | - |

## REFERENCES

Adams, P. D., Afonine, P. V., Bunkóczi, G., Chen, V. B., Davis, I. W., Echols, N., Headd, J. J.,-W. Hung, L., Kapral, G. J., Grosse-Kunstleve, R. W., McCoy, A. J., Moriarty, N. W., Oeffner, R., Read, R. J., Richardson, D. C., Richardson, J. S., Terwilliger, T. C.& Zwart, P. H. (2012). International Tables for Crystallography.539–547.

Afonine, P. V., Poon, B. K., Read, R. J., Sobolev, O. V., Terwilliger, T. C., Urzhumtsev, A. &Adams, P. D. (2018). Acta Crystallogr D Struct Biol.74, 531–544.

Cheng, A., Eng, E. T., Alink, L., Rice, W. J., Jordan, K. D., Kim, L. Y., Potter, C. S. &Carragher, B. (2018). J. Struct. Biol.204, 270–275.

Emsley, P., Lohkamp, B., Scott, W. G.& Cowtan, K. (2010). Acta Crystallogr. D Biol.Crystallogr.66, 486–501.

Glaeser, R. M., Typke, D., Tiemeijer, P. C., Pulokas, J.& Cheng, A. (2011). J. Struct.Biol.174, 1–10.

Henderson, R., Baldwin, J. M., Downing, K. H., Lepault, J.& Zemlin, F. (1986).Ultramicroscopy.19, 147–178.

Herzik, M. A., Jr, Wu, M.& Lander, G. C. (2017).Nat. Methods. 14, 1075–1078.

Kim, L. Y., Rice, W. J., Eng, E. T., Kopylov, M., Cheng, A., Raczkowski, A. M., Jordan, K. D., Bobe, D., Potter, C. S.& Carragher, B.(2018). Front Mol Biosci.5, 50.

Lander, G. C., Stagg, S. M., Voss, N. R., Cheng, A., Fellmann, D., Pulokas, J., Yoshioka, C., Irving, C., Mulder, A., Lau, P.-W., Lyumkis, D., Potter, C. S.& Carragher, B. (2009).Journal of Structural Biology.166, 95–102.

Li, Y., Cash, J. N., Tesmer, J. J. G. & Cianfrocco, M. A. (2019) bioRxiv https://doi.org/10.1101/2019.12.20.885541

Rohou, A. & Grigorieff, N. (2015). J. Struct. Biol.192, 216–221.

Suloway, C., Pulokas, J., Fellmann, D., Cheng, A., Guerra, F., Quispe, J., Stagg, S., Potter, C. S. & Carragher, B. (2005). J. Struct. Biol.151, 41–60.

Wagner, T., Merino, F., Stabrin, M., Moriya, T., Antoni, C., Apelbaum, A., Hagel, P., Sitsel, O., Raisch, T., Prumbaum, D., Quentin, D., Roderer, D., Tacke, S., Siebolds, B., Schubert, E., Shaikh, T. R., Lill, P., Gatsogiannis, C. & Raunser, S. (2019).Commun Biol.2, 218.

Wu, M., Lander, G. C. & Herzik, M. A. (2019) bioRxiv https://doi.org/10.1101/855643.

Zheng, S. Q., Palovcak, E., Armache, J.-P., Verba, K. A., Cheng, Y. & Agard, D. A.(2017). Nat. Methods.14, 331–332.

Zivanov, J., Nakane, T., Forsberg, B. O., Kimanius, D., Hagen, W. J., Lindahl, E. &Scheres, S. H. (2018). Elife.7,.

Zivanov, J., Nakane, T. & Scheres, S. H. W.(2019). IUCrJ.6, 5–17.

Zivanov, J., Nakane, T. & Scheres, S. H. W.(2020). IUCrJ. 7, 253–267.

